# StrainHub: A phylogenetic tool to construct pathogen transmission networks

**DOI:** 10.1101/650283

**Authors:** Adriano de Bernardi Schneider, Colby T. Ford, Reilly Hostager, John Williams, Michael Cioce, Ümit V. Çatalyürek, Joel O. Wertheim, Daniel Janies

## Abstract

In exploring the epidemiology of infectious diseases, networks have been used to reconstruct contacts among individuals and/or populations. Summarizing networks using pathogen metadata (e.g., host species and place of isolation) and a phylogenetic tree is a nascent, alternative approach. In this paper, we introduce a tool for reconstructing transmission networks in arbitrary space from phylogenetic information and metadata. Our goals are to provide a means of deriving new insights and infection control strategies based on the dynamics of the pathogen lineages derived from networks and centrality metrics. We created a web-based application, called StrainHub, in which a user can input a phylogenetic tree based on genetic or other data along with characters derived from metadata using their preferred tree search method. StrainHub generates a transmission network based on character state changes in meta-data, such as place or source of isolation, mapped on the phylogenetic tree. The user has the option to calculate centrality metrics on the nodes including betweenness, closeness, degree, and a new metric, the source/hub ratio. The outputs include the network with values for metrics on its nodes and the tree with characters reconstructed. All of these results can be exported for further analysis.

**Availability:** https://github.com/abschneider/StrainHub and strainhub.io

## Introduction

New technologies can shape responses to outbreaks of rapidly evolving infectious diseases. High-throughput genetic sequencing has allowed rapid characterization of disease outbreaks such as Ebola, Yellow Fever, and Zika viruses (1–3). Multiple advancements in interpreting genomic data related to pathogen outbreaks have been recently developed: SCOTTI (4), Phyloscanner (5), QUENTIN (6), BadTrIP (7), Outbreaker (8) and Outbreaker2 (9). Each of these tools offer distinct advantages when analyzing datasets, although only a few of them include network visualization and use centrality metrics in order to infer importance of nodes (6).

Here, we introduce a novel tool, StrainHub, which summarizes the transition between states of metadata rather than among individuals, providing an overview of the pathogen transmission paths through geography or populations. Genomic data and associated metadata from pathogens are observations of the biology underlying a disease. The combination of these data collected from related pathogen isolates allow researchers to understand disease transmission patterns as a function of the pathogens’ evolutionary history. StrainHub can leverage these data and make phylogenetic transmission graphs accessible to public health scientists. StrainHub is provided as both a standalone package in GitHub and a web-based interface.

## Methods

We built the StrainHub application with multiple R packages wrapped with Shiny. In order to build a transmission network, the user provides a phylogenetic tree and associated metadata for all terminal taxa.

### Ancestry reconstruction

When parsimony is selected, the ancestral state reconstruction step in StrainHub uses the R function “asr_max_parsimony” from the package Castor (10). This function performs an ancestral state reconstruction for discrete traits derived from metadata using the parsimony algorithm described by Sankoff (11). Next, based on the results of ancestral state reconstruction, StrainHub outputs a relationship list of the metadata elements as source and destination.

Users have the option to run phylogeography in BEAST (12) and visualize the transmission network based on the tree edges and trait probability for each node. The relationship list is extracted from the tree nodes of a BEAST phylogeography file using the R package treeio (13) to build a directional network and calculate the metrics as described below. This option is used in lieu of the parsimony ancestral state reconstruction step.

### Tree and transmission network visualization

We implemented a tree visualization tab using ggtree, ggplot, and plotly to display the metadata mapped to each taxon within the tree (14, 15). We used the R packages igraph and visNetwork to build the transmission networks. igraph provides functions for generating the backbone of the transmission network and calculates the centrality metrics (16). visNetwork provides the R interface to the “vis.js” javascript charting library, allowing an interactive visualization of the transmission network (17). StrainHub uses the edge list created on the previous step as a source and destination list which is transformed into a data frame and plotted as a network. The nodes of the transmission network created are not the individual pathogen sequences but the relationship of the ancestral and descendant states of the pathogen sequences (e.g., changes in geography, host shifts, changes among risk factors, or any set of discrete state the user can encode).

### Transmission network metrics

In StrainHub we provide multiple centrality measurements to determine the relative importance of nodes within a network with respect to the dynamics of pathogen lineages (18, 19). In this application, we implemented three centrality metrics: betweenness, closeness, and degree (Table 1). Betweenness measures the number of shortest paths between two other nodes that pass through the node of interest, normalized by the number of all pairs of nodes within the network. The higher betweenness score of a node reflects the importance of that node as a hub in the network for traffic of the pathogen. Closeness evaluates a node based on the relative sum of the lengths of all the shortest paths from that node to all other nodes within network. A higher closeness centrality value is associated with how close a node is as a direct point of transmission to other nodes. Degree is the number of edges incident upon a given node. The higher the number of edges connected to a given node indicates the higher importance of that node in terms of being involved in metadata state transitions irrespective of directionality (20).

**Table 1.**
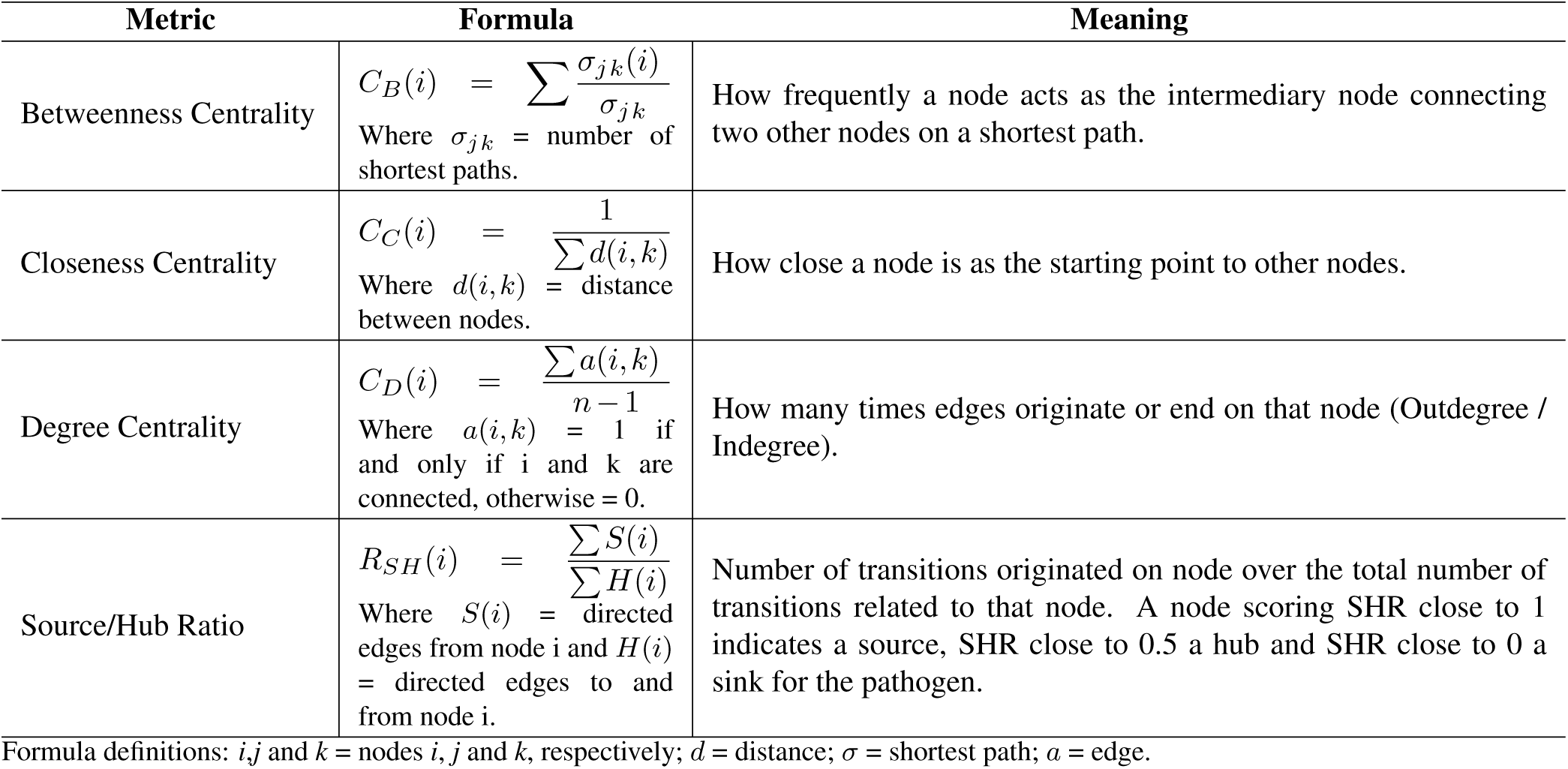
Summary of Centrality Metrics applied to measure transmission networks on StrainHub.

We further divided degree centrality into “indegree” and “outdegree”. With these values, we created the “Source Hub Ratio” (SHR) metric. SHR is obtained by calculating the ratio of all transitions from the node (outdegree) over all transitions from and to the node (indegree + outdegree), resulting in a value ranging from 0 to 1. Nodes with SHR values close to 0.5 indicate that the node is a hub, SHR close to 1, indicate it is a source, and SHR close to 0 indicates that it is a sink for the pathogen. The SHR metric reflects the importance of a node within the network as the hub, source or sink of the pathogen, ignoring centrality of the node within network (21). Although SHR alone does not define which node is the most important within the network for the spread of the pathogen, it creates an indicator for the behavior of the nodes, which can be further investigated for importance by the association of SHR with other metrics of interested for the user. The availability of multiple metrics for any type of discrete metadata allows the user to have flexibility in assessing hypotheses in different contexts for the spread of infectious diseases.

### Shiny

We used the Shiny framework to provide StrainHub a flexible, web-based interface. Shiny allows for the generation of interactive applications that can be hosted as standalone web-applications that are locally installed or that are served over a network (22). In ether case, the user interacts with StrainHub via a web browser of choice.

## Implementation

StrainHub accepts phylogenetic trees in Newick format and metadata in comma-separated value (CSV) format to run the ancestry reconstruction step. The orthography and content of the taxon names must match. Alternatively, users can run phylogeography in BEAST and use the maximum clade credibility tree in Nexus format as input to visualize the transmission network.

Three outputs are generated during the transmission network analysis: a transmission network plot, a tree plot with the character of interest mapped to the tips of the tree, and a table with all centrality metrics computed. The transmission network graph and the tree files are user-interactive, and a snapshot for further analysis or publication can be exported in PNG format (e.g., Figure 1). The centrality metrics table can be exported as a CSV file.

**Fig. 1.**
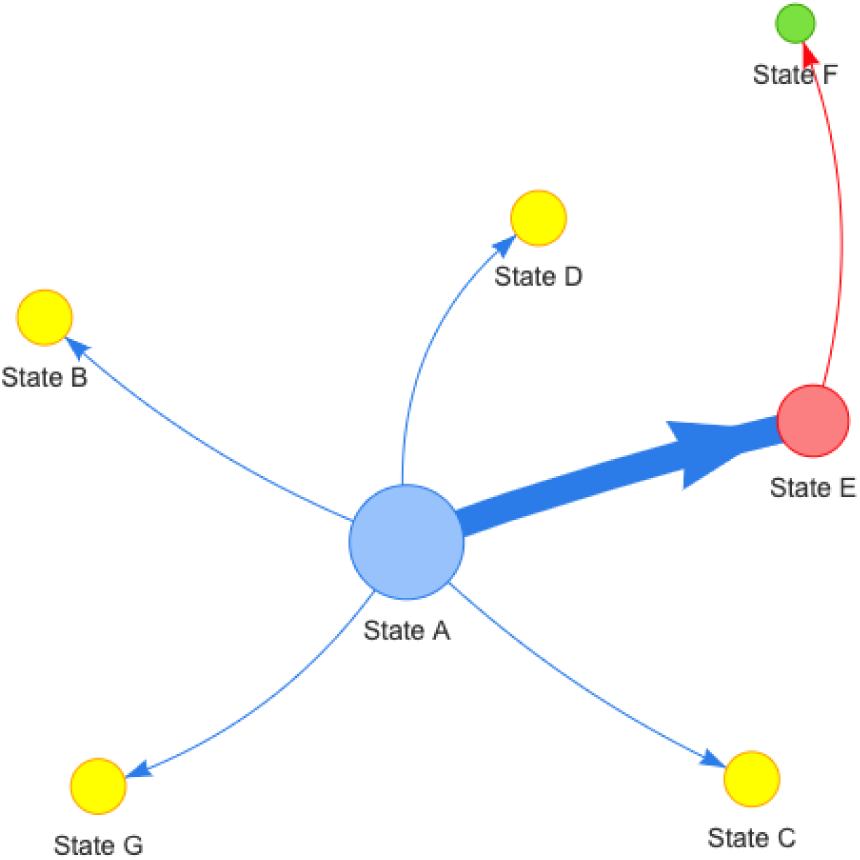
Example of a generic pathogen transmission network based on geographic location (states). Size of nodes are scaled by the metric selected by the user (i.e. Betweenness, Closeness, Degree or Source Hub Ratio), arrows reflect directionality of transition between states, thickness of lines and arrows represents frequency of transitions (thicker arrows = more transitions), colors of nodes are randomly assigned to metric value and associated with nodes with same values.

The web-application can be accessed at strainhub.io and the source code is available at https://github.com/abschneider/StrainHub under the GNU General Public License (GPL) v3.0.

## Conclusion

We created a visual analytic tool that enables the user to interpret and communicate phylogenetic data on the dynamics of the spread pathogens over geography or various hosts. Moreover, the metadata format can be used for any type of categorical data that user can encode (e.g., food sources, or risk factor, or phenotype). The user use should assume the results are only as strong as the underlying phylogenetic data (i.e. sampling across metadata states and taxa, adequate branch lengths and nodal support to reconstruct an ancestor descendent change). With solid datasets, the data in the transmission networks, such as identification of the hubs and sources for the spread of pathogens, will assist health authorities to allocating resources to parts of the network that will do the most to disrupt the spread of the pathogen.

## Acknowledgements

We thank Andrew Frick, Dr. Nídia Trovão, and Dr. Tetyana Vasylyeva for suggestions on methodology during the development of this app.

## Funding

This work was supported in part by the National Institutes of Health (NIH) National Institute of Allergy and Infectious Diseases (grant numbers K01AI110181 and AI135992) to JOW. We acknowledge the support of the Department of Bioinformatics and Genomics, the College of Computing and Informatics, and the Graduate School of the University of North Carolina at Charlotte. This effort was funded in part by the Defense Threat Reduction Agency under contract HDTRA1-16-C-0010 to UC and DJ.

## Notes

#### Summary of Updates

New figure, fixed description of centrality metrics, added to introduction description of other software in the field, author affiliations updated.

http://strainhub.io

https://github.com/abschneider/StrainHub

